# Organization of orbitofrontal-auditory pathways in the Mongolian gerbil

**DOI:** 10.1101/2023.04.25.538296

**Authors:** Rose Ying, Lashaka Hamlette, Laudan Nikoobakht, Rakshita Balaji, Nicole Miko, Melissa L. Caras

**Affiliations:** Neuroscience and Cognitive Science Program, University of Maryland, College Park, Maryland, 20742; Department of Biology, University of Maryland, College Park, Maryland, 20742; Center for Comparative and Evolutionary Biology of Hearing, University of Maryland, College Park, Maryland, 20742

**Author notes:** Correspondence: Rose Ying, Biology-Psychology Building 4094 Campus Dr College Park, MD 20742.

## Abstract

Sound perception is highly malleable, rapidly adjusting to the acoustic environment and behavioral demands. This flexibility is the result of ongoing changes in auditory cortical activity driven by fluctuations in attention, arousal, or prior expectations. Recent work suggests that the orbitofrontal cortex (OFC) may mediate some of these rapid changes, but the anatomical connections between the OFC and the auditory system are not well-characterized. Here, we used virally-mediated fluorescent tracers to map the projection from OFC to the auditory midbrain, thalamus, and cortex in a classic animal model for auditory research, the Mongolian gerbil (*Meriones unguiculatus*). We observed no connectivity between the OFC and the auditory midbrain, and an extremely sparse connection between the dorsolateral OFC and higher-order auditory thalamic regions. In contrast, we observed a robust connection between the ventral and medial subdivisions of the OFC and the auditory cortex, with a clear bias for secondary auditory cortical regions. OFC axon terminals were found in all auditory cortical lamina but were significantly more concentrated in the infragranular layers. Tissue-clearing and lightsheet microscopy further revealed that auditory cortical-projecting OFC neurons send extensive axon collaterals throughout the brain, targeting both sensory and non-sensory regions involved in learning, decision-making, and memory. These findings provide a more detailed map of orbitofrontal-auditory connections and shed light on the possible role of the OFC in supporting auditory cognition.

## Introduction

Sensory perception is critical for survival, enabling organisms to find food and mates, evade predators, and navigate to safety. To perform these behaviors successfully, individuals must parse a seemingly endless array of complex time-varying sensory stimuli and respond appropriately. Accordingly, perceptual sensitivity is highly malleable, shifting from moment to moment as a result of fluctuations in attentional state (Ahissar & Hochstein, 1993; Luck & Ford, 1998; Shinn-Cunningham, 2008; Spitzer et al., 1988; Yotsumoto & Watanabe, 2008), arousal level (Asutay & Västfjäll, 2012; Kim et al., 2017; Mather & Sutherland, 2011; McGinley et al., 2015), and expectations (Ashton, 2014; Carcea et al., 2017; de Lange et al., 2018; Pinto et al., 2015; Stein & Peelen, 2015; Vangkilde et al., 2012). These dynamics allow organisms to filter out irrelevant information, make rapid predictions about upcoming sensory events, and more easily detect, discriminate, or identify signals of high interest or value.

Perceptual flexibility results from rapid changes in stimulus-driven activity within sensory cortices. Reductions in response variability (McGinley et al., 2015; von Trapp et al., 2016), adaptive transformations of receptive field properties (Atiani et al., 2009; David et al., 2012; Fontanini & Katz, 2006; Fritz et al., 2003; Fritz et al., 2005, 2007; Yin et al., 2014), reorganizations of neuronal networks (Cohen & Maunsell, 2009; Downer et al., 2017; Francis et al., 2018, 2022; Sheikhattar et al., 2018; Steinmetz et al., 2000), and shifts in response gain (Caras & Sanes, 2017; Chapman & Meftah, 2005; Davis, 1964; Hubel et al., 1959; Niwa et al., 2012; Spitzer et al., 1988; Treue & Maunsell, 1999; Yoshida & Katz, 2011) have all been reported. Many of these rapid changes are thought to be mediated by descending inputs from frontal cortical brain regions to sensory cortices. While significant attention has been devoted to studying the contribution of medial prefrontal cortical areas (Bahmani et al., 2019; Shinn-Cunningham, 2017), a growing body of work suggests that the orbitofrontal cortex (OFC) also plays an important role. OFC is unique in that it is the only prefrontal cortical region to receive input from all sensory modalities (Le Merre et al., 2021), suggesting its central involvement in sensory processing. More recent work has revealed its role in the top-down control of sensory cortical function (Banerjee et al., 2020; Liu et al., 2020; Liu et al., 2021).

The OFC’s contribution to auditory cognition is particularly evident. OFC neurons respond to sounds (Sharma & Bandyopadhyay, 2020; Srivastava & Bandyopadhyay, 2020; Winkowski et al., 2013) and in humans, the magnitude of OFC activity predicts aspects of auditory perception and behavior. For example, increased OFC gain is associated with phantom sound perception during tinnitus (Lee et al., 2020; Vanneste et al., 2012). Conversely, OFC lesions impair tone recognition and weaken auditory cortical responsiveness (Kam et al., 2018). This latter finding, which links OFC damage to a reduction in auditory cortical activity, is consistent with studies in mice that demonstrate that OFC activation can alter sound-evoked representations in the primary auditory cortex. Specifically, pairing sounds with electrical stimulation of the OFC (Winkowski et al., 2013) or with optogenetic stimulation of OFC axon terminals in the auditory cortex (Winkowski et al., 2018) shifts the frequency tuning of auditory cortical neurons and alters auditory cortical network structure. These changes improve the ability of the auditory cortical neuronal population to discriminate the paired sound frequency.

While connections between the OFC and auditory regions have been identified in several species, including house mice (*Mus musculus*) (Sharma & Bandyopadhyay, 2020; Srivastava & Bandyopadhyay, 2020; Winkowski et al., 2018; Zingg et al., 2014), Lister-hooded rats (*Rattus norvegicus*) (Olthof et al., 2019), rhesus macaque monkeys (*Macaca mulatta*) (Cavada et al., 2000; Hackett et al., 1999; Romanski, Bates, et al., 1999; Romanski, Tian, et al., 1999), and humans (Cammoun et al., 2015), we lack a detailed characterization of these connections in a classic and well-established model for auditory research, the Mongolian gerbil (*Meriones unguiculatus*). Gerbils offer several unique advantages for auditory studies. Unlike other commonly used rodent models, like mice or rats, gerbils exhibit excellent low-frequency hearing capabilities, similar to humans (Ryan, 1976). Additionally, there is a large body of literature on gerbil auditory anatomy (Budinger et al., 2000a, 2000b, 2008; Budinger & Scheich, 2009; Radtke-Schuller et al., 2016), development (Anbuhl et al., 2022; Caras & Sanes, 2015, 2019; Müller, 1996; Rosen et al., 2010; Sanes, 1993; Sanes & Rubel, 1988), sound perception (Jüchter et al., 2022; von Trapp et al., 2016; Yao et al., 2020), auditory skill learning (Caras & Sanes, 2017, 2019; Sarro & Sanes, 2011; Wetzel et al., 1998), hearing loss (Henry et al., 1980; Mills et al., 1990; Takesian et al., 2012; Tucci et al., 1999; von Trapp et al., 2017; Xu et al., 2007), and central nervous system function (Amaro et al., 2021; Belliveau et al., 2014; Buran et al., 2014; Franken et al., 2018; Kreeger et al., 2021; Lesica et al., 2010; Myoga et al., 2014; Yao & Sanes, 2021), spanning multiple decades. A more complete description of the inputs from the OFC to the gerbil auditory system would facilitate our understanding of the neural circuits that support rapid adjustments to the acoustic environment, and may additionally shed light on the mechanistic link between hearing status and cognitive function (Lin et al., 2011; Taljaard et al., 2016).

Here, we used virally mediated fluorescent tracers to characterize the descending projections from the OFC to the gerbil auditory cortex, thalamus, and midbrain. We report a heterogeneous pattern of auditory connectivity across the mediolateral axis of the OFC. Dorsolateral OFC (DLO) sends a sparse projection to the auditory thalamus and exhibits little to no connectivity with the auditory midbrain or cortex. In contrast, ventral (VO) and medial (MO) subdivisions of the OFC innervate primary auditory cortex (Au1) as well as the secondary dorsal (AuD) and ventral (AuV) cortices, with the densest labeling in the deep infragranular layers. Further investigation using tissue clearing and lightsheet microscopy revealed that auditory cortical-projection neurons also send collaterals to distant targets, including non-auditory sensory cortices and brain regions implicated in learning, decision-making, and memory. Together, our findings reveal the organization of orbitofrontal-auditory connectivity in the gerbil brain and provide additional anatomical context for the OFC’s contribution to auditory cognition.

## Materials and methods

### Animals

Mongolian gerbils were obtained from Charles River and bred in-house. Gerbils were group housed on a 12-hour light cycle and given *ad libitum* access to chow (Purina Mills Lab Diet 5001) and water. All procedures except one were conducted at the University of Maryland College Park, and were approved by the University of Maryland Institutional Animal Care and Use Committee. The remaining single procedure (subject ID_269953, see Table 1) was conducted at New York University and was approved by the New York University Institutional Animal Care and Use Committee.

**Table 1.**
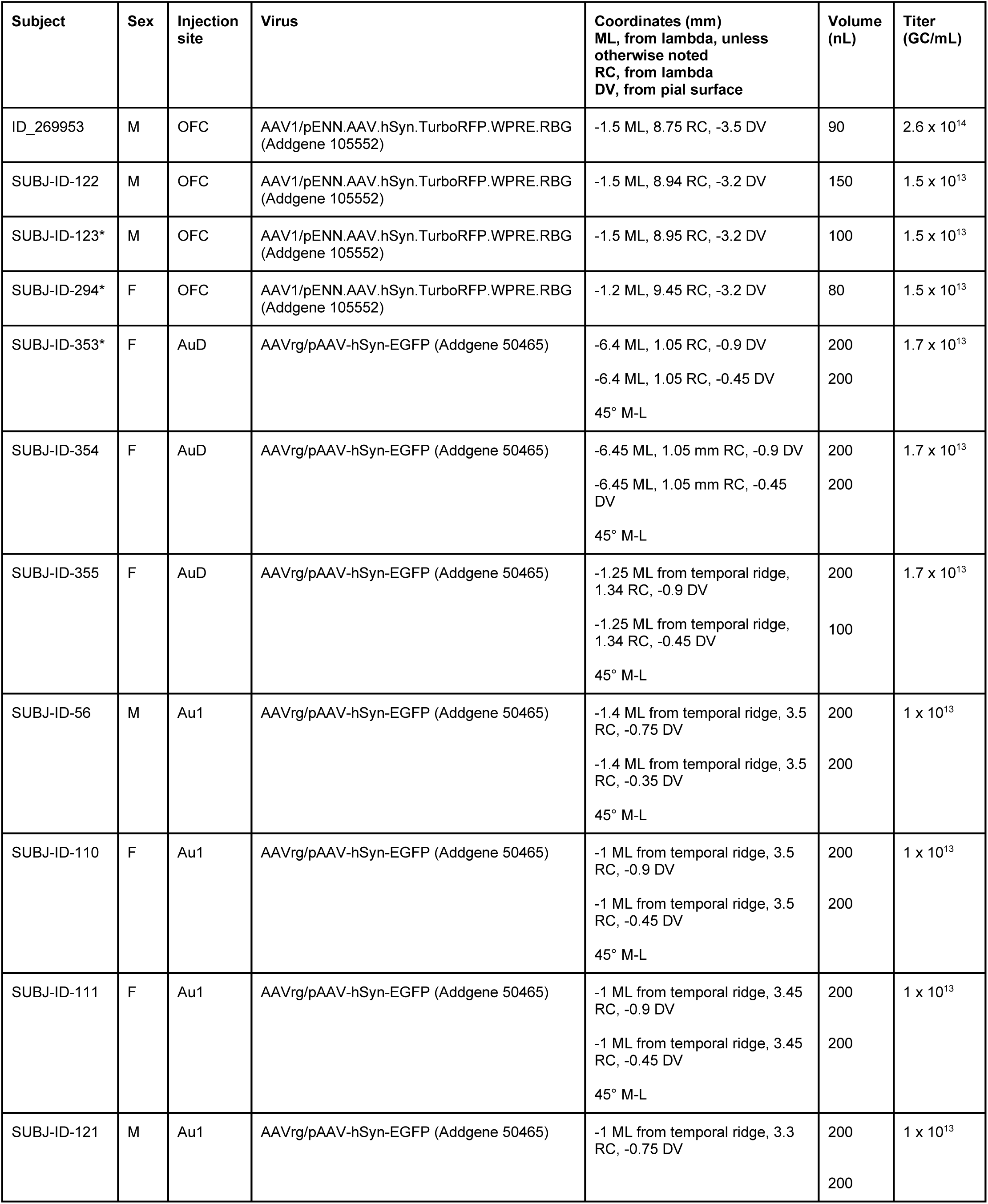
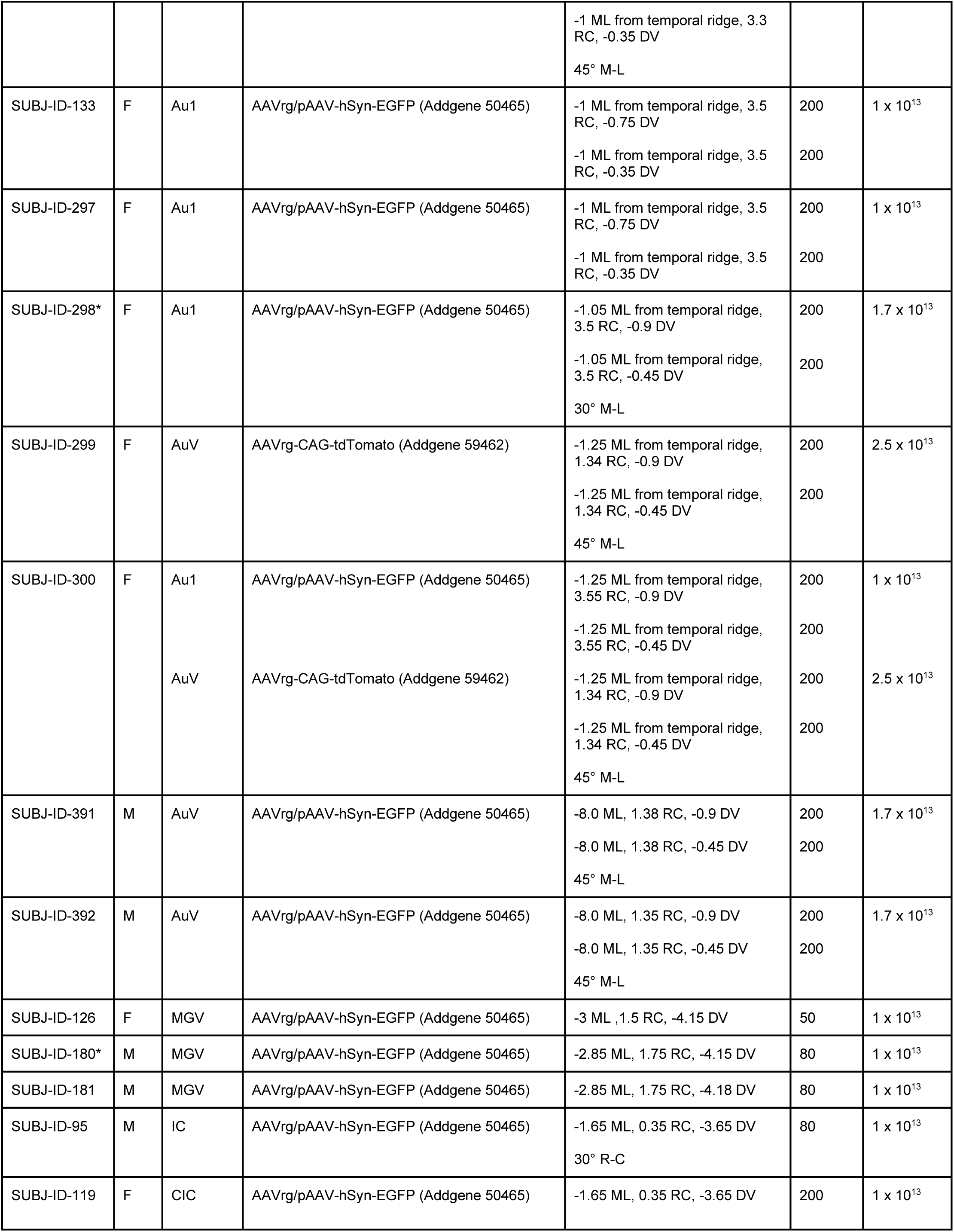
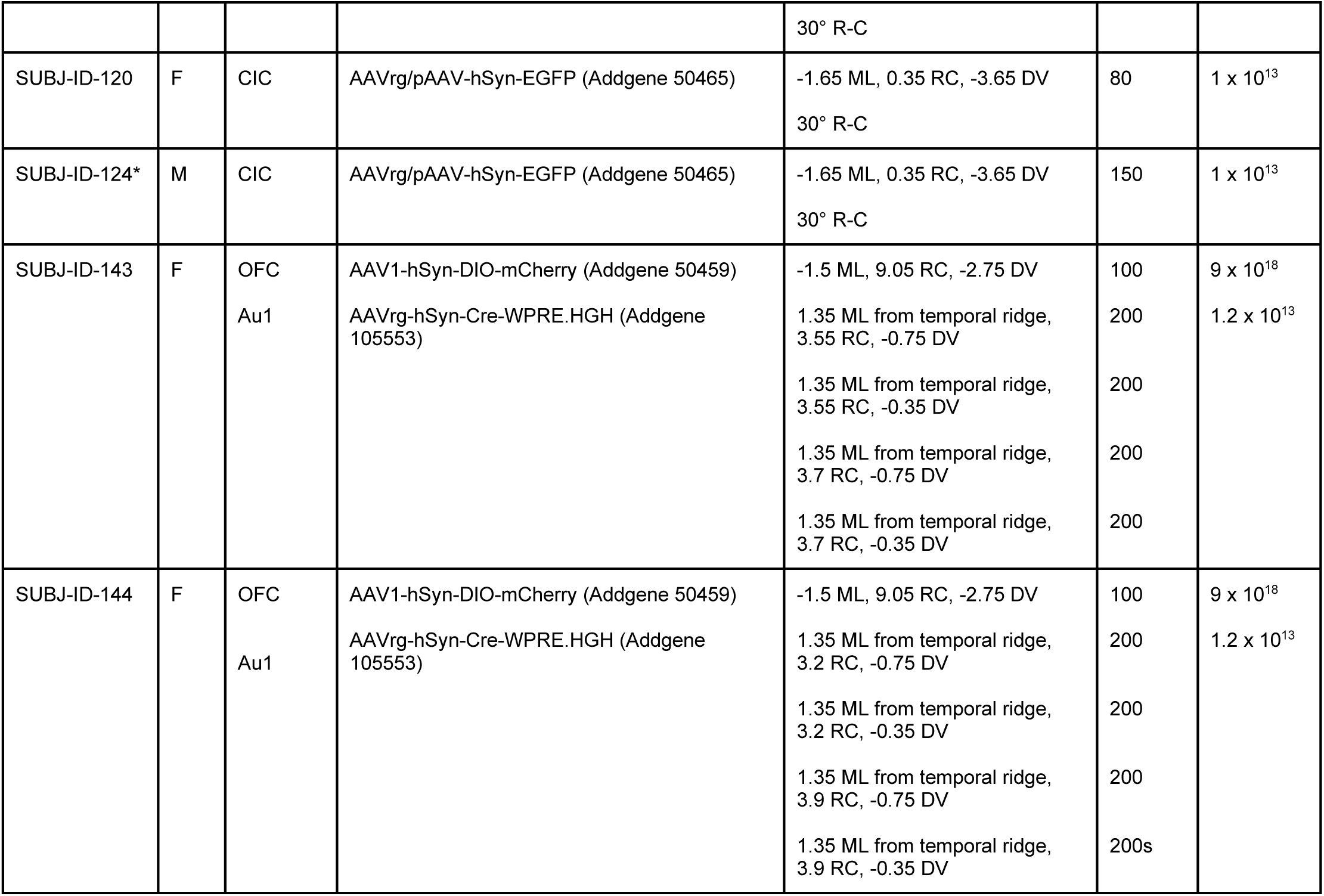
Subjects and injection coordinates. Pipette angle = 0° unless otherwise noted. * denotes representative subject used for figure.

### Viral injections

Subjects were injected with one or more viral tracers into the OFC, or into subcortical or cortical auditory regions. For complete details about each injection, including the virus, injection volume, titer, and coordinates used, see Table 1. A total of 44 animals (aged postnatal day 58-637) were used in this study. The data from off-target injections (N = 10), injections with excessive virus spread outside the region of interest (N = 5), and brains with excessive tissue damage (N = 2) were excluded from all analyses. Therefore, this report includes data from 27 animals (N = 16 female).

One day prior to surgery, animals were given meloxicam (1.5 mg/kg, 1.5 mg/mL) either via oral suspension or subcutaneous injection or a carprofen medigel cup as a preventative analgesic. On the day of surgery, animals were administered meloxicam or carprofen and dexamethasone (0.35 mg/kg, 0.5 mg/mL) subcutaneously to prevent edema. Subjects were placed in a small induction chamber in which isoflurane (5%) was continuously administered in O_2_ (2 L/min). After the animal was sedated, the fur on the animal’s head was shaved. The animal was then transferred to a warming pad on a stereotaxic device (Kopf) and secured in place with ear bars and a bite bar. A nose cone delivered a steady flow of isoflurane (1.5-2.5%) and oxygen at a flow rate of 2 L/min. Once a surgical plane of anesthesia was achieved (as evidenced by slow and steady respiration, and the absence of a toe-pinch response) ophthalmic ointment was applied to the eyes, and the surgical area was treated with alternating applications of betadine and alcohol. A midline incision was made, and the skin and fascia were retracted. The skull was cleaned and dried with H_2_O_2_ and a craniotomy was made over the region of interest using a 5mm drill bit. A glass injection pipette was filled with mineral oil and attached to a programmable injector (Nanoject III, Drummond). The virus was drawn up into the tip and positioned over the injection site. A durotomy was performed, and the pipette was inserted to the targeted depth. The virus was injected at a rate of 2 nL/sec and allowed to diffuse for 5-15 mins prior to pipette retraction. The craniotomy was covered with KwikSil and the skin was sutured along the incision. Immediately after the surgery, subjects were given a subcutaneous injection of Normosol (1-2 mL) and allowed to recover in their home cage. An additional dose of meloxicam or carprofen and dexamethasone were administered 24 hours after surgery, and subjects were closely monitored after the surgical procedure.

### Histology

Subjects remained in their home cages for 3-5 weeks after injections to allow for protein expression, after which animals were perfused for histology. Animals were anesthetized with an intraperitoneal injection of ketamine (150 mg/kg, 25 mg/mL) and xylazine (6 mg/kg, 1 mg/mL) or sodium pentobarbital (60-100 mg/kg, 50 mg/mL) in saline. After the subject was determined to be unresponsive to a toe pinch, the chest cavity was opened and the subject was perfused transcardially using 1x phosphate buffered saline (PBS) followed by 4% paraformaldehyde.

### Tissue processing and confocal microscopy

Extracted brains were post-fixed in 4% paraformaldehyde at 4°C for at least one day, then embedded in 6% agar and sliced on a vibratome (Leica VT1000 S) at 70 μm thickness. Slices were mounted on gelatin subbed slides and dried overnight. Mounted sections were counterstained with NeuroTrace 435/455 Fluorescent Nissl (Invitrogen N21479). The slices were rehydrated with 1x PBS for at least 30 minutes. After a 5-minute wash in 1x PBS, the slices were incubated in 1x PBS with 0.1% Triton X for 20 minutes. The slides were washed three times for 5 minutes each in 1x PBS, then incubated in a 1:100 NeuroTrace dilution overnight at 4°C. After incubation, the slices were washed for 10 minutes in 1x PBS with 0.1% triton, then twice in 1x PBS for 5 minutes each. The slides were then cover-slipped using ProLong Diamond Antifade Mountant and imaged using a confocal microscope (Zeiss LSM 980). Post-processing, including stitching and z-stack processing, were completed using Zeiss ZEN and FIJI software (Schindelin et al., 2012).

### Tissue clearing and lightsheet microscopy

For lightsheet imaging experiments, fixed brains were extracted, packaged in 1x PBS with 0.02% sodium azide and sent to LifeCanvas Technologies in Cambridge, MA. Brains were cleared using SHIELD and the mCherry signal was enhanced via immunolabeling in SmartBatch+ (7.5 μg rabbit anti-RFP). Cleared whole brains were imaged using a lightsheet microscope at 3.6X magnification. Regions of particular interest were imaged again at 15X magnification. Clearing and image acquisition was performed by LifeCanvas Technologies.

### Cell body quantification in OFC

Brain sections separated by 250-300μm along the rostral to caudal dimension were imaged. Each image was matched to a section in the brain atlas (Radtke-Schuller et al., 2016). Anterior and posterior OFC were defined by the emergence of the claustrum, consistent with previous reports in rodents (Barreiros et al., 2021a; Barreiros et al., 2021b). Manual cell counts were performed for each image, and cell counts were independently verified by another lab member. Cell counts were normalized to enable statistical comparisons between hemispheres and between subregions within hemispheres. For complete details about cell counts and corresponding plate numbers in the gerbil brain atlas, see Table 2.

**Table 2.**
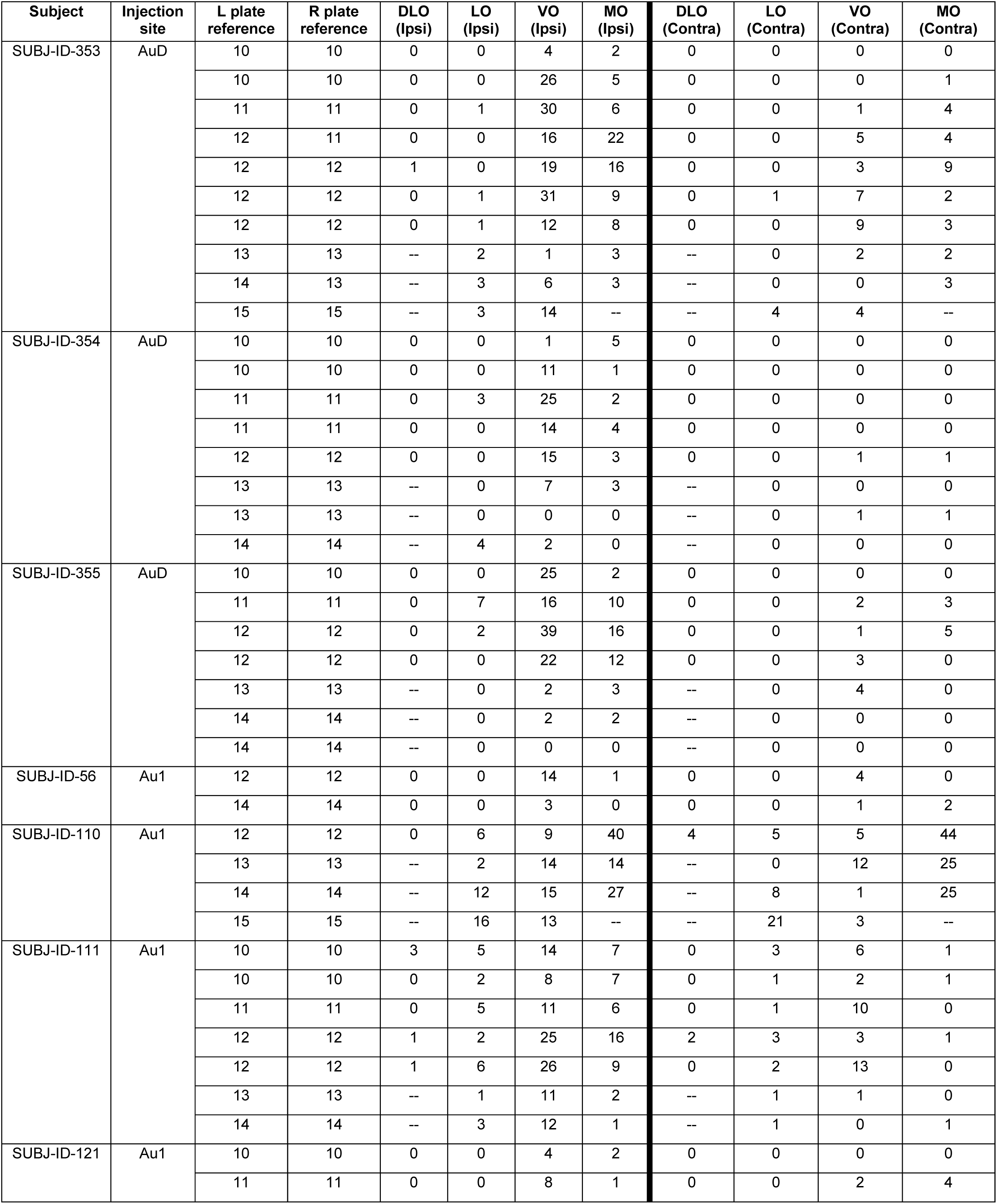
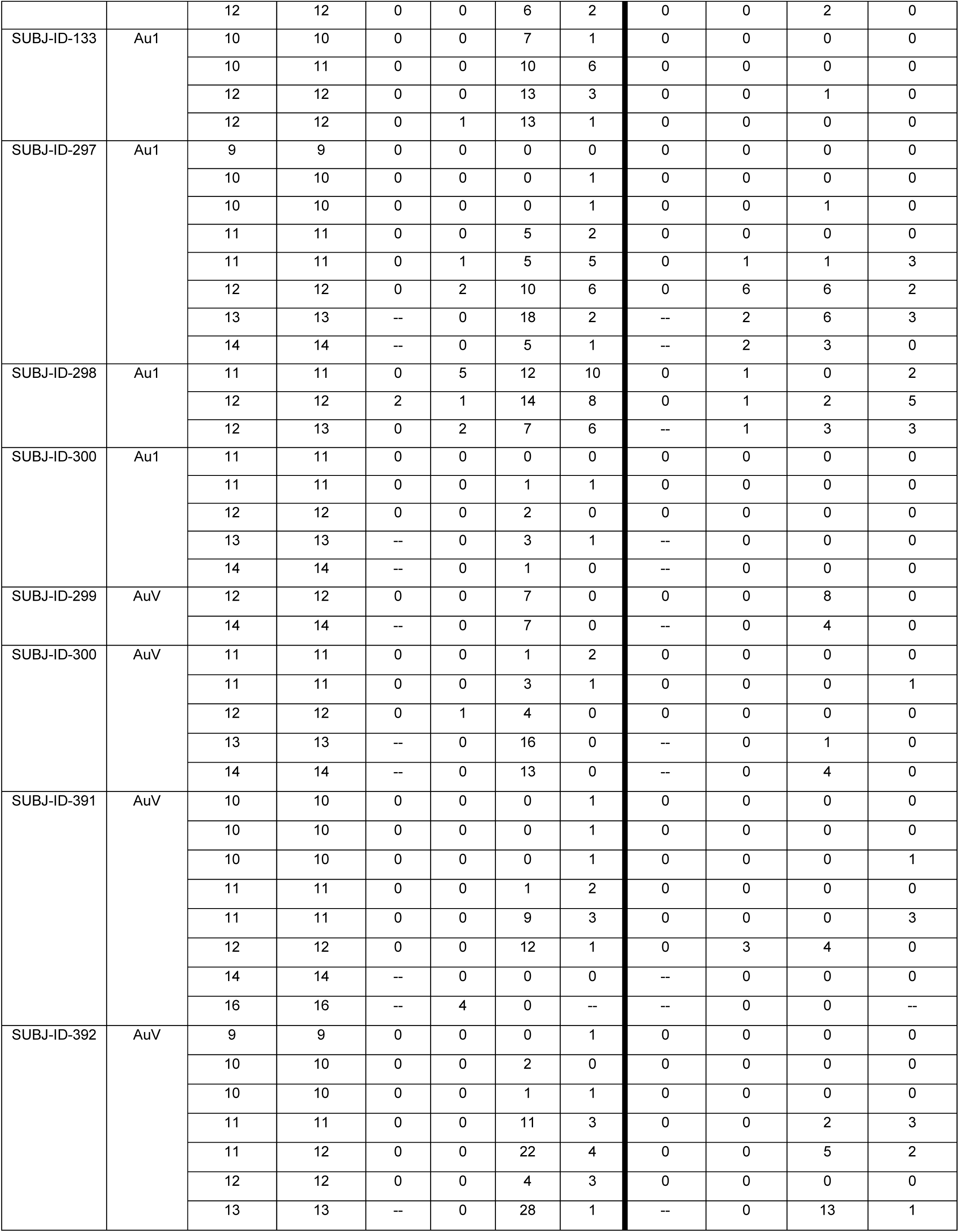
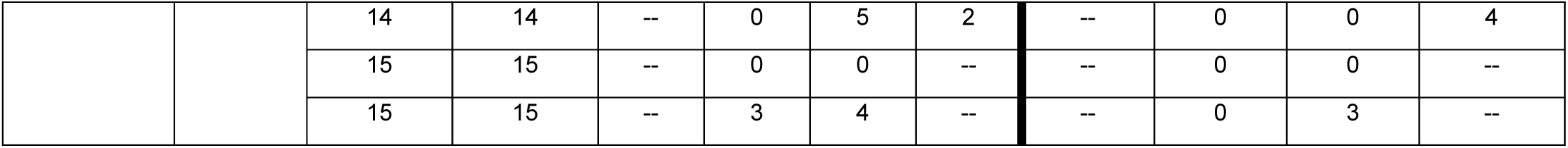
Raw OFC cell count data. Plate reference refers to the plate number in the gerbil brain atlas (Radtke-Schuller et al., 2016).

### Fiber analysis in the auditory cortex

Intensity analysis was performed using FIJI. For each image, color channels were split into red, green, and blue, and the appropriate channel was converted to grayscale (red for turboRFP, green for EGFP). Excess background was removed. The regions of interest were identified by comparing the Nissl stain and other gross morphological features to the gerbil brain atlas (Radtke-Schuller et al., 2016) and marked on the image. Intensity values were calculated using the Plot Profile tool along linear transects, drawn from the cortical surface through the cortical layers at roughly equal distances through the region of interest. Custom MATLAB code was written to bin intensity values (200 bins per sample) as a function of transect length. Intensity values were then normalized to the maximum intensity value for each image. For each subject, one image was taken in the anterior portion of the auditory cortex (plates 28-30 in the gerbil brain atlas) and one image from the posterior portion (plates 31-33). To compare between anterior and posterior sections, each set of anterior and posterior images was processed so that the intensity histogram range, median, and standard deviation between images were as similar as possible.

### Axon collateral analysis

Lightsheet images were reconstructed into a 3-D model in Imaris Viewer. The location of labeled fibers, their apparent density, and the presence of boutons were qualitatively described.

### Statistical analysis

For analysis of cell bodies in the OFC, generalized linear mixed models (GLMs) and paired t-tests were performed in JMP Pro 15. Subject and subregion were included as nested categorical random effects (subregion within subject). To quantify the intensity of fiber labeling throughout the auditory cortex, we used R to construct a GLM using the ‘glmmTMB’ (Brooks et al., 2017), DHARMa (Hartig, 2022), car (Fox & Weisberg, 2018), and emmeans (Lenth, 2023) packages. Subject, region (AuD/Au1/AuV), layer (1-6), and position (anterior/posterior) were included as categorical random effects and quadruple nested (position within layer within region within subject). Normal distribution of residuals was tested after running the GLM and an analysis of variance (ANOVA) was used to analyze the statistical significance of the GLM. Post-hoc pairwise comparisons were conducted after transforming data back to the original scale.

We considered results statistically significant when p < 0.05. All p values were adjusted for multiple comparisons using a Bonferroni correction and were corrected for any violations of sphericity using the Greenhouse-Geisser method.

## Results

### Retrograde tracing from core auditory cortex

A previous study reported the presence of retrogradely labeled cell bodies in the OFC after small injections of fluorescent dextran tracers were delivered to gerbil Au1, but the distribution pattern of labeled soma within the OFC was not reported (Budinger et al., 2008). Rodent OFC is classically divided into distinct subregions along its mediolateral axis, including the dorsolateral/agranular insular (DLO/AI), lateral (LO), ventral (VO), and medial (MO) areas (Krettek & Price, 1977; Öngür & Price, 2000; Price, 2006; Ray & Price, 1992; Rempel-Clower, 2007). These subdivisions exhibit different patterns of anatomical connectivity, and are thought to mediate distinct behavioral and cognitive functions (for review see Barreiros et al., 2021a; Bradfield & Hart, 2020; Izquierdo, 2017; Rempel-Clower, 2007). These observations raise the question of whether the OFC neurons that innervate the auditory cortex are evenly distributed across the mediolateral axis, or primarily originate in one or more discrete subregions.

To answer this question, we injected AAVrg-hSyn-EGFP, a viral vector in a serotype that allows for robust retrograde access to long-range projection neurons (Tervo et al., 2016), into left core auditory cortex. Correctly placed injections (N = 8; see Table 1) were centered in Au1 and/or the anterior auditory field (AAF) and spanned multiple cortical layers (Figure 1A, B). In two cases, the injection site exhibited minimal upward spread, just beyond the border with secondary somatosensory cortex (S2) and AuD; in the remaining cases the injections were contained entirely within the boundaries of Au1/AAF.

**Figure 1.**
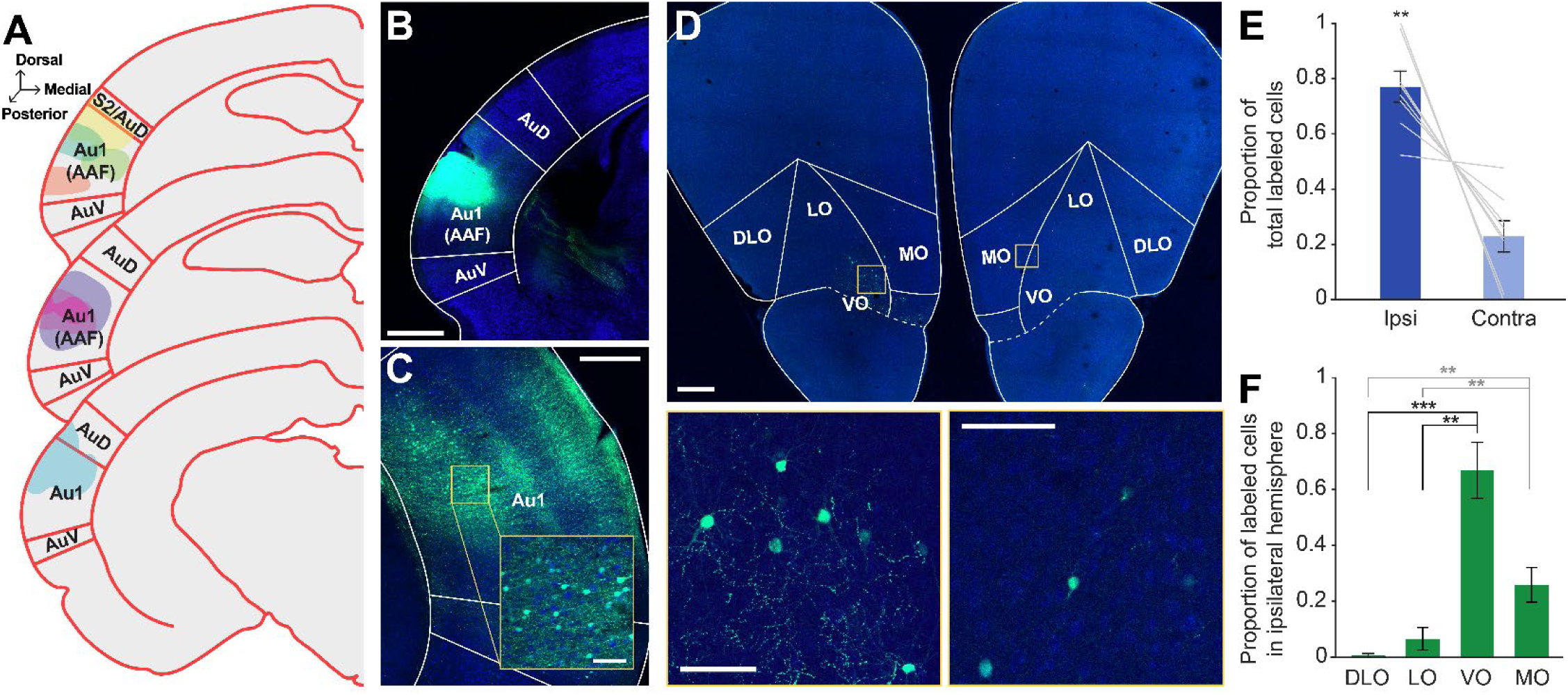
Retrograde tracer into Au1 reveals distinct profile of cell body labeling across the OFC mediolateral axis. **A.** Schematic of all injection sites in Au1 (N = 8). **B.** Injection site of representative subject. Scale bar = 1000 μm. **C.** In the same subject as B, expected labeling observed in the contralateral Au1. Scale bar = 500 μm, inset = 100 μm. **D.** Representative OFC slice in the same subject as B. High magnification images depict cell bodies in the ipsilateral (left) and contralateral (right) hemispheres. Scale bar = 500 μm, high-mag = 100 μm. **E.** Proportion of total labeled cells in each hemisphere. Bars represent means ± standard error; lines connect data points from individual animals. **F.** Proportion of cell bodies within the ipsilateral hemisphere localized to each OFC subregion. Bars represent means ± standard error. **p<0.01, ***p<0.001.

To first confirm that our retrograde tracing approach worked as expected, we took advantage of the known interhemispheric connection between auditory cortices (Budinger et al., 2000a; Diamond et al., 1968; Imig & Reale, 1980), and examined the contralateral auditory cortex for the presence of labeled cell bodies. As expected, we observed many brightly labeled neurons throughout the contralateral auditory cortex (Figure 1C). In the OFC, we observed clearly labeled cell bodies scattered throughout the region (Figure 1D). A significantly higher proportion of the labeled neuron population was in the ipsilateral hemisphere compared to the contralateral hemisphere (t_7_ = −4.80, p = 0.002) (Figure 1E). Given that the proportion of contralateral labeled neurons was significantly smaller, we focused our subsequent analyses on the ipsilateral hemisphere only. Within the ipsilateral hemisphere, cell body labeling was unevenly distributed across OFC subregions (GLM, F_3,28_ = 45.31, p < 0.0001; Figure 1F). The highest proportion of labeled cell bodies was consistently observed in the VO, followed by MO. Very little labeling was evident in LO or DLO/AI. A complete statistical breakdown of these results is provided in Table 3.

**Table 3.**
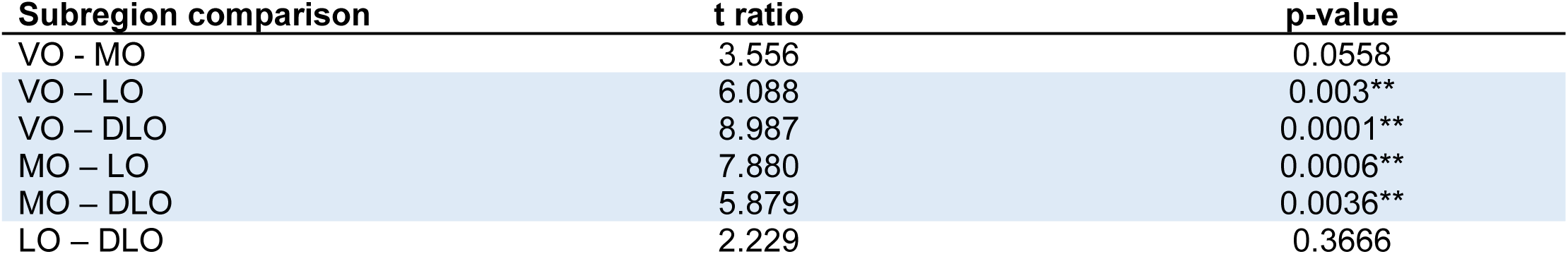
Paired t-tests comparing proportion of labeled cell bodies in each OFC subregion after retrograde injections into Au1 (see Figure 1). All p-values were corrected for multiple comparisons using a Bonferroni adjustment.

| Subregion comparison | t ratio | p-value |
| --- | --- | --- |
| VO - MO | 3.556 | 0.0558 |
| VO – LO | 6.088 | 0.003** |
| VO – DLO | 8.987 | 0.0001** |
| MO – LO | 7.880 | 0.0006** |
| MO – DLO | 5.879 | 0.0036** |
| LO – DLO | 2.229 | 0.3666 |

### Retrograde tracing from secondary auditory cortices

Non-sensory processes, like attention and behavioral context, modulate the activity of higher-order auditory regions, and the magnitude of these modulations are often larger than those observed in Au1 (Atiani et al., 2014; Jäncke et al., 1999; Niwa et al., 2015; O’Sullivan et al., 2019; Yin et al., 2020). This finding suggests that non-sensory brain regions may differentially connect to primary and secondary auditory cortices. We therefore asked whether OFC neurons project to two secondary auditory cortical regions in the gerbil—the dorsal (AuD) and ventral (AuV) auditory cortices.

First, we injected a retrograde tracer (AAVrg-hSyn-EGFP) into the left AuD. Well-placed injections (N = 3) spanned most or all cortical layers and exhibited little to no spread beyond the boundaries of AuD (Figure 2A, B). As expected, we identified cell bodies in the contralateral AuD, confirming the efficacy of retrograde expression (Figure 2C) (Budinger et al., 2000a). An examination of OFC revealed brightly labeled cell bodies densely clustered in the ventral portion of the region (Figure 2D). Similar to Au1-projecting neurons, the vast majority of AuD-projecting neurons were located in the ipsilateral hemisphere (t_2_ = 7.81, p = 0.016; Figure 2E).

**Figure 2.**
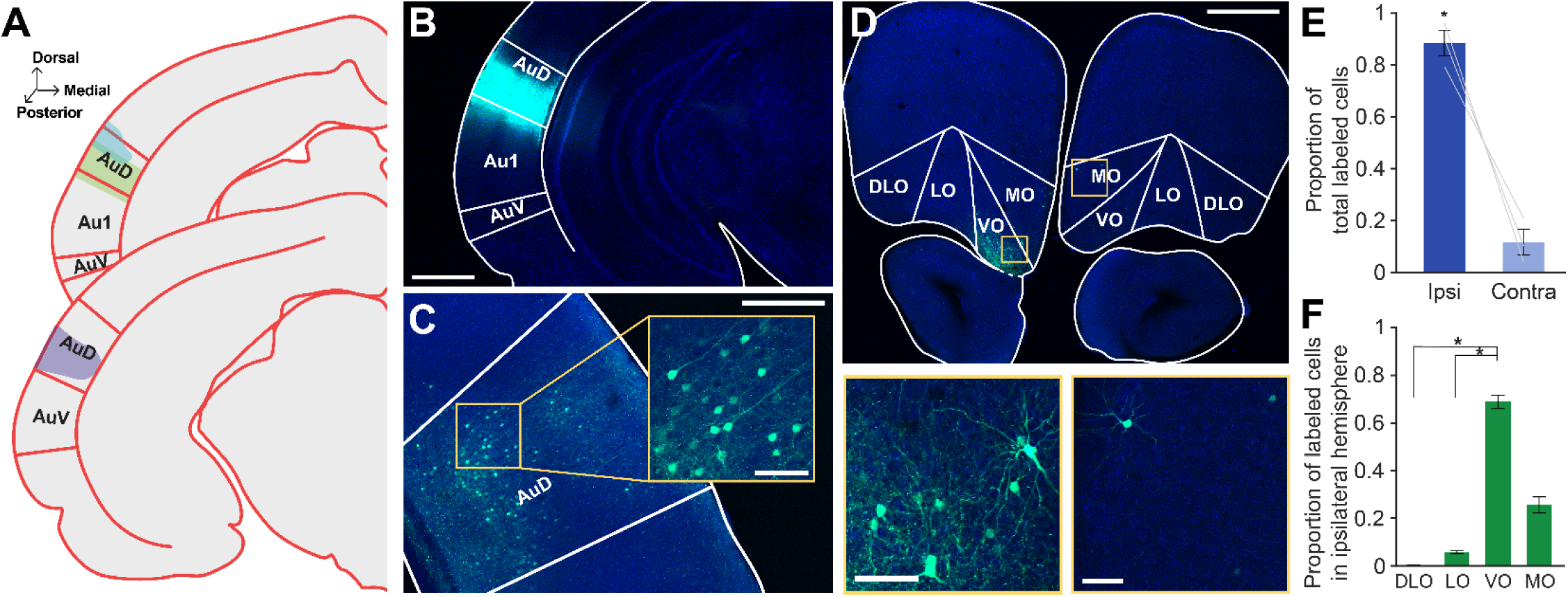
Retrograde tracer into AuD labels cell bodies in OFC. **A.** Schematic of all injection sites in left AuD (N = 3). **B.** Injection site of representative subject. Scale bar = 1000 μm. **C.** In the same subject as in B, expected labeling observed in contralateral AuD. Scale bar = 500 μm, inset = 100 μm. **D.** Labeled OFC cell bodies in the same subject as B. High magnification images depict cell bodies in the ipsilateral (left) and contralateral (right) OFC. Scale bar = 500 μm, high-mag = 100 μm. **E.** Proportion of total labeled cells in each hemisphere. Bars represent means ± standard error; lines connect data points from individual animals. **F.** Proportion of cell bodies within the ipsilateral hemisphere localized to each OFC subregion. Bars represent means ± standard error. *p<0.05

We then quantified apparent differences in cell body labeling across the mediolateral axis of the ipsilateral OFC. Labeled neurons were unequally distributed across OFC subregions (GLM, F_3,8_ = 655.64, p < 0.0001), and the pattern was similar to that observed for Au1/AAF-projecting neurons: VO contained the highest proportion of labeled cell bodies, followed by MO, LO, and DLO/AI (Figure 2F). A complete statistical breakdown of these results is provided in Table 4.

**Table 4.**
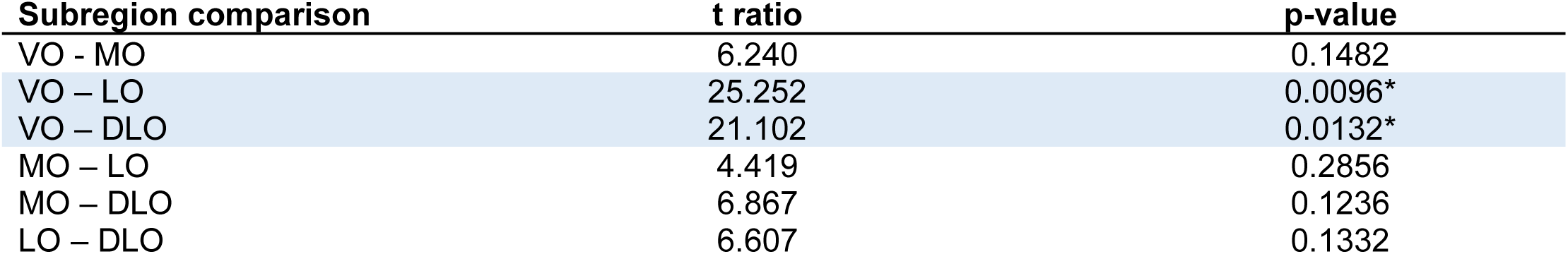
Paired t-tests comparing proportion of labeled cell bodies in each OFC subregion after retrograde injection into AuD (see Figure 2). All p-values were corrected for multiple comparisons using a Bonferroni adjustment.

| Subregion comparison | t ratio | p-value |
| --- | --- | --- |
| VO - MO | 6.240 | 0.1482 |
| VO – LO | 25.252 | 0.0096* |
| VO – DLO | 21.102 | 0.0132* |
| MO – LO | 4.419 | 0.2856 |
| MO – DLO | 6.867 | 0.1236 |
| LO – DLO | 6.607 | 0.1332 |

We also injected retrograde tracers (AAVrg-hSyn-EGFP or AAVrg-CAG-tdTomato) into the AuV. Injections (N = 4) were centered in AuV, with one injection exhibiting minor spread upwards into AuD (Figure 3A, B). We identified cell bodies in the contralateral AuV to confirm the efficacy of the retrograde virus (Figure 3C) (Budinger et al., 2000a). As in Au1 and AuD, cell bodies were present in the OFC, with a significantly higher proportion of cells on the ipsilateral side (t_2_ = 3.29, p = 0.047; Figure 3D, E). Across the mediolateral axis, we once again observed unequal distribution of labeled neurons (GLM, F_3,12_ = 62.35, p < 0.0001; Figure 3F). VO contained the highest proportion of labeled cells, followed by MO, LO, and DLO. A complete statistical breakdown of these results is provided in Table 5.

**Figure 3.**
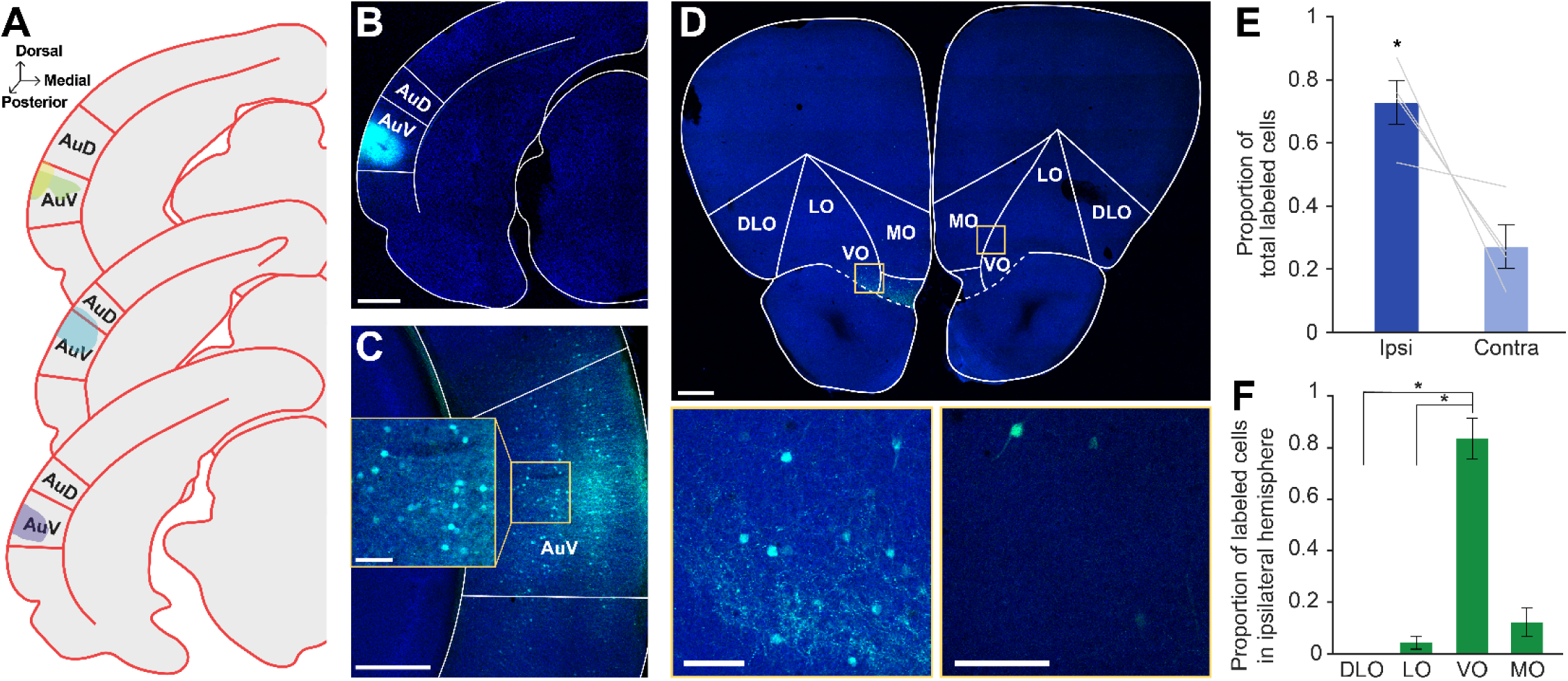
Retrograde tracer into AuV labels cell bodies in OFC. **A.** Schematic of all injection sites in left AuV (N = 4). **B.** Injection site of representative subject. Scale bar = 1000 μm. **C.** In the same subject as in B, expected labeling observed in contralateral AuV. Scale bar = 500 μm, inset = 100 μm. **D.** Labeled OFC cell bodies in the same subject as B. High magnification images depict cell bodies in the ipsilateral (left) and contralateral (right) OFC. Scale bar = 500 μm, insets = 100 μm. **E.** Proportion of total labeled cells in each hemisphere. Bars represent means ± standard error; lines connect data points from individual animals. **F.** Proportion of cell bodies within the ipsilateral hemisphere localized to each OFC subregion. Bars represent means ± standard error. *p<0.05

**Table 5.**
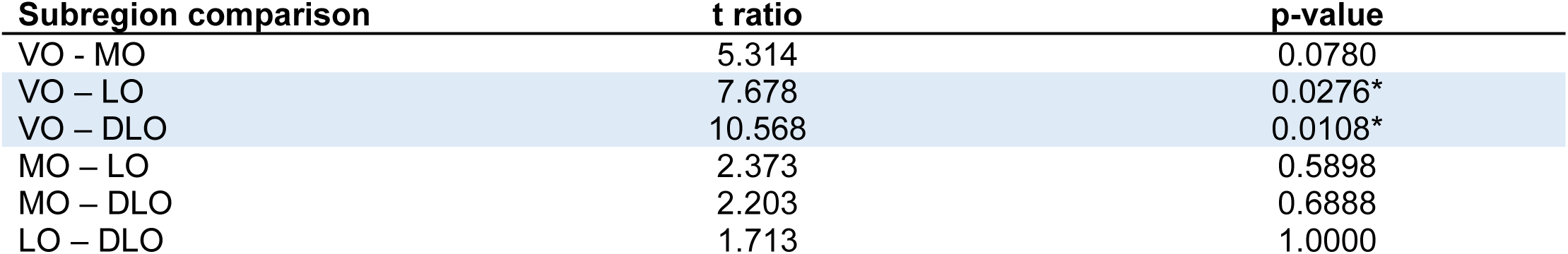
Paired t-tests comparing proportion of labeled cell bodies in each OFC subregion after retrograde injection into AuV (see Figure 3). All p-values were corrected for multiple comparisons using a Bonferroni adjustment.

Qualitatively, we observed that more OFC cell bodies were present after retrograde tracing from secondary auditory cortices than from Au1 (e.g., compare Figures 1D, 2D, and 3D). These results were not quantified due to uneven sample sizes across groups and variability in injection spread across cortical layers.

### Retrograde tracing from the auditory thalamus and midbrain

Subcortical auditory regions, including the medial geniculate nucleus (MGN) and inferior colliculus (IC), also play important roles in sound processing, cognition, and multi-modal integration (Bartlett & Wang, 2007; Brainard & Knudsen, 1993; Ehret & Merzenich, 1988; Wang et al., 2008). Both of these structures exhibit context-dependent modulations of sound-evoked responses (Franceschi & Barkat, 2021; Gilad et al., 2020; Gruters & Groh, 2012; Mihai et al., 2021; Ryan et al., 1984; Ryan & Miller, 1977; Slee & David, 2015; von Kriegstein et al., 2008), and the activation of prefrontal cortex shapes neural activity in the MGN (Barry et al., 2017). These findings suggest that direct inputs from frontal cortical areas may influence subcortical auditory processing. In line with this idea, one previous study reported that neurons in the ventral and lateral OFC innervate the central nucleus of the rat IC (Olthof et al., 2019). These observations led us to ask whether a similar connection exists between the OFC and the gerbil IC and/or MGN.

Towards that end, we injected a retrograde tracer (AAVrg-hSyn-eGFP) into the left MGN of three subjects. Viral expression was primarily localized to the ventral subdivision of the MGN (MGV), which serves as the primary lemniscal input to Au1 (Budinger et al., 2000b), but exhibited some spread into the nearby non-lemniscal dorsal (MGD) subdivision and the medial zone of the medial geniculate (MZMG) (Figure 4A,B). To confirm retrograde expression, we documented labeled cell bodies in the IC, which provides the majority of ascending input to the MGN (Cant & Benson, 2007; Hackett, 2011; Kudo & Niimi, 1978; Ledoux et al., 1987; Morest, 1965) (Figure 4C).

**Figure 4.**
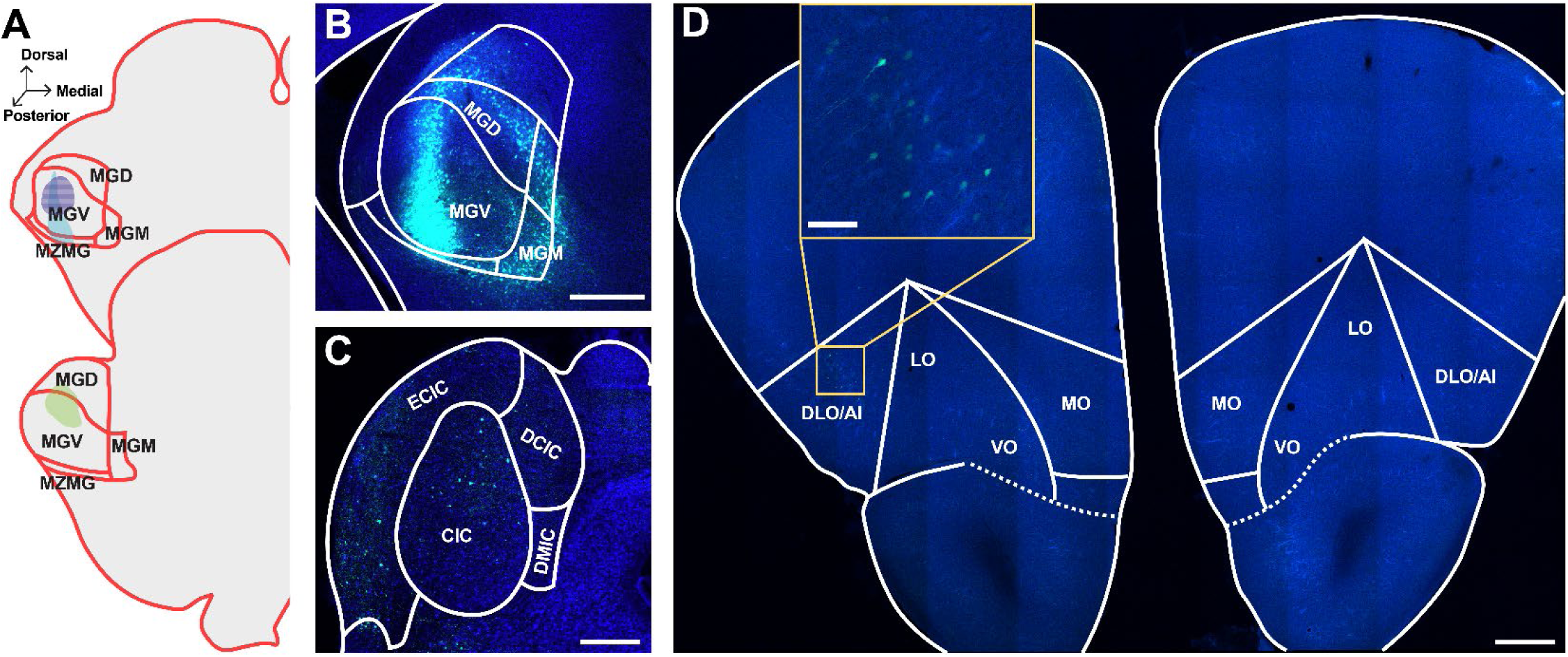
Retrograde tracer into MGN sparsely labels cell bodies in OFC. **A.** Schematic of all injection sites in MGN (N = 3). Striped pattern denotes an injection that resulted in no observed cell body labeling in the OFC. **B.** Injection site of representative subject. Scale bar = 500 μm. **C.** In the same subject as B, expected labeling observed in IC. Scale bar = 500 μm. **D.** Representative OFC slice in the same subject as B. Scale bar = 500 μm, inset = 100 μm.

Two of the three MGN injections resulted in sparse cell body labeling in the OFC, primarily in the ipsilateral DLO/AI (Figure 4D). The remaining injection did not result in any OFC cell body labeling. Notably, this injection was entirely confined to the MGV, with no spread into non-leminscal areas (see purple striped injection pattern in Figure 4A). These data suggest that MGN-projecting neurons in the OFC primarily terminate in higher-order auditory thalamic subdivisions. Because these injections resulted in relatively few labeled cell bodies in the OFC, we did not quantify these findings.

To investigate whether OFC projects to IC, we injected four animals with AAVrg-hSyn-eGFP into the left IC. All injections were centered in the central nucleus of the IC (CIC), which serves as the primary ascending input to the MGV (Hackett, 2011; Morest, 1965), with moderate spread into the overlying external and dorsal IC cortices (Figure 5A,B). As expected from decades of previous work characterizing a prominent corticofugal pathway from Au1 to IC (Bajo et al., 2010; Bajo & Moore, 2005; Budinger et al., 2000b; Cooper & Young, 1976; Games & Winer, 1988; Suga et al., 2000; Williamson & Polley, 2019) we observed distinct cell body labeling in layer 5 of Au1 (Figure 5C). Despite this confirmation of successful retrograde cortical expression, and in contrast to an earlier report in rat (Olthof et al., 2019), no cell bodies were observed in any of the OFC subregions in either hemisphere (Figure 5D).

**Figure 5.**
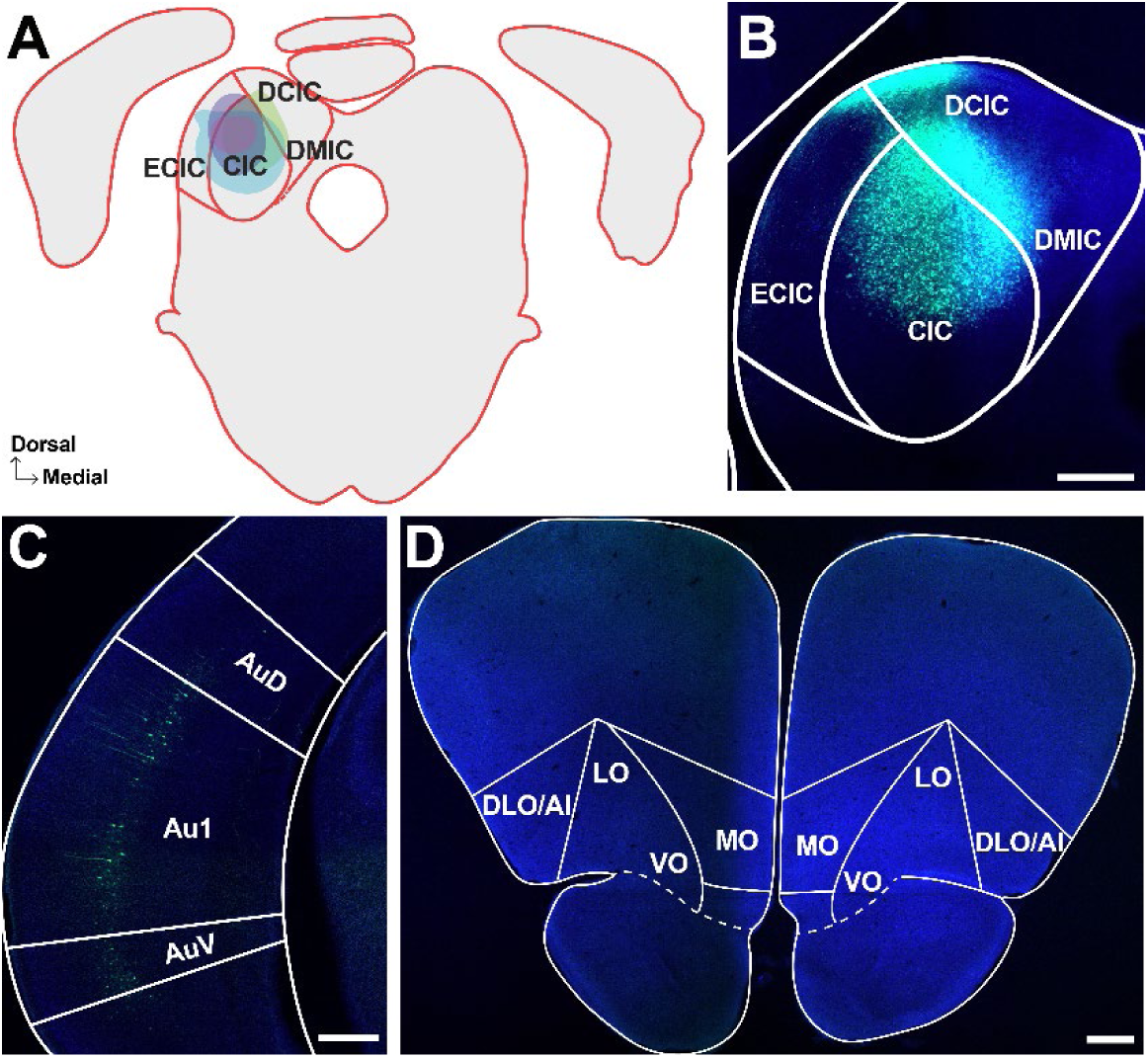
Retrograde tracer into IC does not label cell bodies in OFC. **A.** Schematic of all injection sites in IC (N = 3). **B.** Injection site of representative subject. Scale bar = 500 μm. **C.** Expected labeling observed in Au1. Scale bar = 500 μm. **D.** Representative OFC slice in the same subject as B showing no cell bodies. Scale bar = 500 μm.

### Anterograde tracing from OFC

Context-dependent changes in sound-evoked activity exhibit a distinct laminar profile in the auditory cortex (Atencio & Schreiner, 2010; Franceschi & Barkat, 2021; Francis et al., 2018; O’Connell et al., 2014; Sugimoto et al., 1997). Determining the pattern of OFC innervation across auditory cortical layers could therefore provide insight into the potential functional role of the OFC in guiding sound-driven behavior. We injected AAV1-hSyn-TurboRFP into the left OFC of four animals to confirm our retrograde tracing results and to characterize the translaminar pattern of OFC axon distribution in the gerbil auditory cortex. Well-placed injections were centered within the boundaries of OFC (generally LO and VO), with minimal upward spread into the prelimbic and motor cortices and/or downward spread into the piriform cortex (Figure 6A, B).

**Figure 6.**
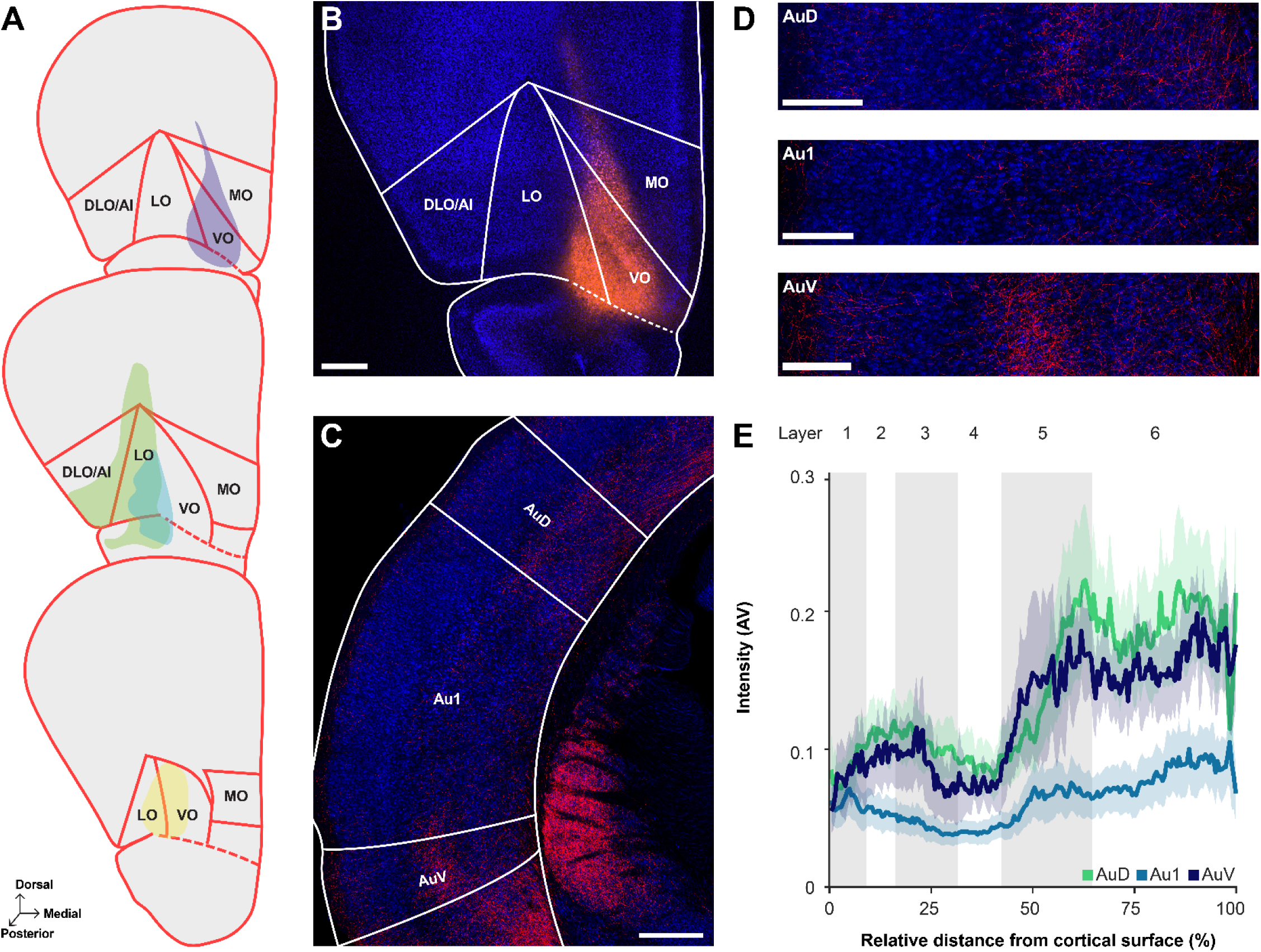
Anterograde tracer into OFC reveals region- and layer-specific innervation pattern in auditory cortex. **A.** Schematic of all injection sites in OFC (N = 4). **B.** Injection site of representative subject. Scale bar = 500 μm. **C.** Labeled terminals in the auditory cortex. Scale bar = 500 μm. **D.** From the same subject as in C, close-up images of terminals through AuD (top), Au1 (middle), and AuV (bottom). Scale bars = 250 μm. **E.** Normalized label intensity (arbitrary value, AV) for each subregion as a function of relative distance from the cortical surface. Lines represent means across animals; shading represents standard error.

As shown in Figure 6C, brightly labeled fibers were observed in both primary (Au1) and secondary (AuD and AuV) auditory cortices, and the labeling intensity differed significantly across these regions (GLM/ANOVA, χ^2^(2) = 22.87, p < 0.0001) (Figure 6D-E). Post-hoc tests revealed that overall label intensity was similar in AuD and AuV (t_103_ = 0.517, p = 0.8632), and significantly stronger in both regions compared to Au1 (AuD: t_103_ = 3.514, p = 0.0019; AuV: t_103_ = 3.206, p = 0.005). These findings indicate that OFC neurons preferentially target higher-order auditory cortical areas over Au1, consistent with our qualitative retrograde tracing observations. Fiber labeling intensity significantly differed overall across cortical layers (GLM/ANOVA, χ^2^(5) = 119.0933, p < 0.0001; Figure 6D-E). Post-hoc tests showed that in contrast to the relatively sparse fibers observed in superficial and granular layers, the infragranular layers (5 and 6) exhibited robust labeling. A full statistical breakdown of these results is provided in Table 6. There was no significant difference in terminal intensity along the anterior-posterior axis (GLM/ANOVA, χ^2^(1) = 0.3890, p = 0.5328; not shown).

**Table 6.**
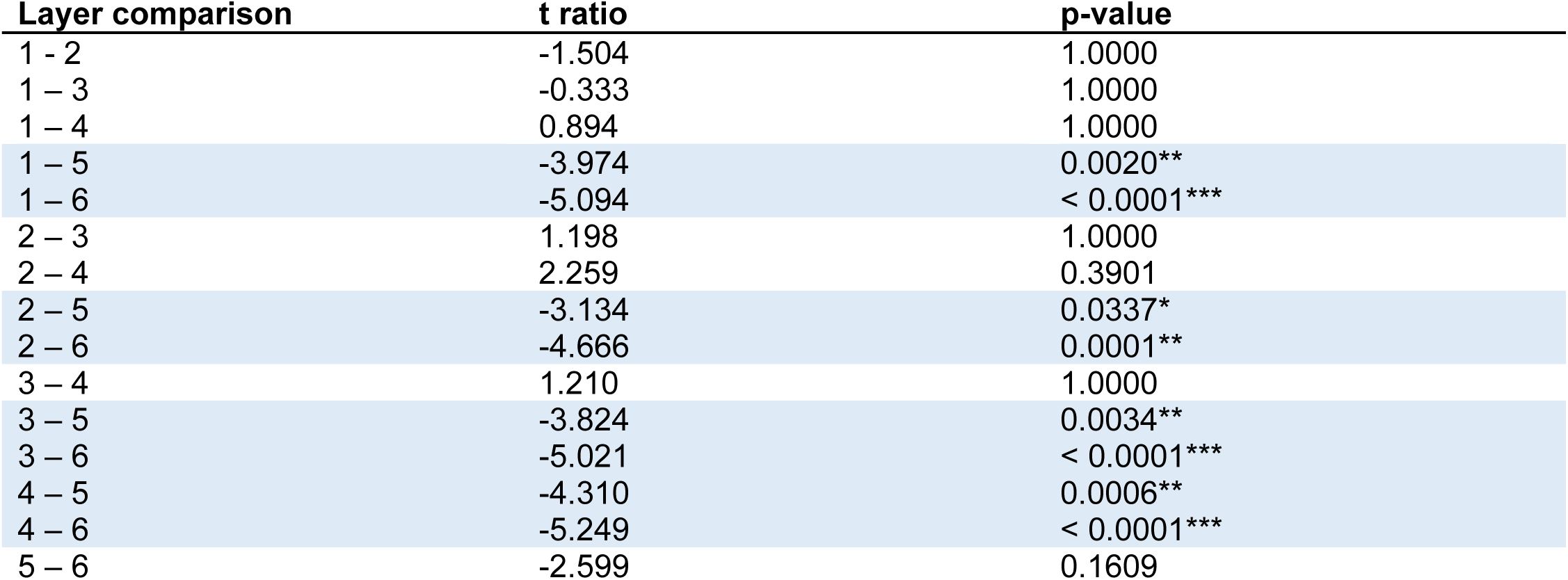
Paired t-tests comparing intensity of labeled OFC axon fibers across auditory cortical layers (see Figure 6). All p values were corrected for multiple comparisons using a Bonferroni adjustment.

| Layer comparison | t ratio | p-value |
| --- | --- | --- |
| 1 - 2 | -1.504 | 1.0000 |
| 1 - 3 | -0.333 | 1.0000 |
| 1 - 4 | 0.894 | 1.0000 |
| 1 - 5 | -3.974 | 0.0020** |
| 1 - 6 | -5.094 | < 0.0001*** |
| 2 - 3 | 1.198 | 1.0000 |
| 2 - 4 | 2.259 | 0.3901 |
| 2 - 5 | -3.134 | 0.0337* |
| 2 - 6 | -4.666 | 0.0001** |
| 3 - 4 | 1.210 | 1.0000 |
| 3 - 5 | -3.824 | 0.0034** |
| 3 - 6 | -5.021 | < 0.0001*** |
| 4 - 5 | -4.310 | 0.0006** |
| 4 - 6 | -5.249 | < 0.0001*** |
| 5 - 6 | -2.599 | 0.1609 |

Our retrograde tracing experiments revealed a sparse connection between the OFC and MGN (Figure 4) and no connection between OFC and IC (Figure 5). To determine whether these observations are genuine, or whether they instead reflect a limited ability of the AAV-retro serotype to access corticothalamic and corticocollicular projections (Tervo et al. 2016), we asked whether anterogradely labeled axon fibers were present in the MGN and/or IC of subjects injected with AAV1-hSyn-TurboRFP into the OFC. In line with our retrograde findings, three injections resulted in sparse labeling throughout the MGN (Figure 7A), with the fourth animal excluded as images were not taken of subcortical auditory areas in this subject. These findings confirm that OFC does in fact innervate MGN, albeit weakly. No clear labelling was observed in the IC (Figure 7B).

**Figure 7.**
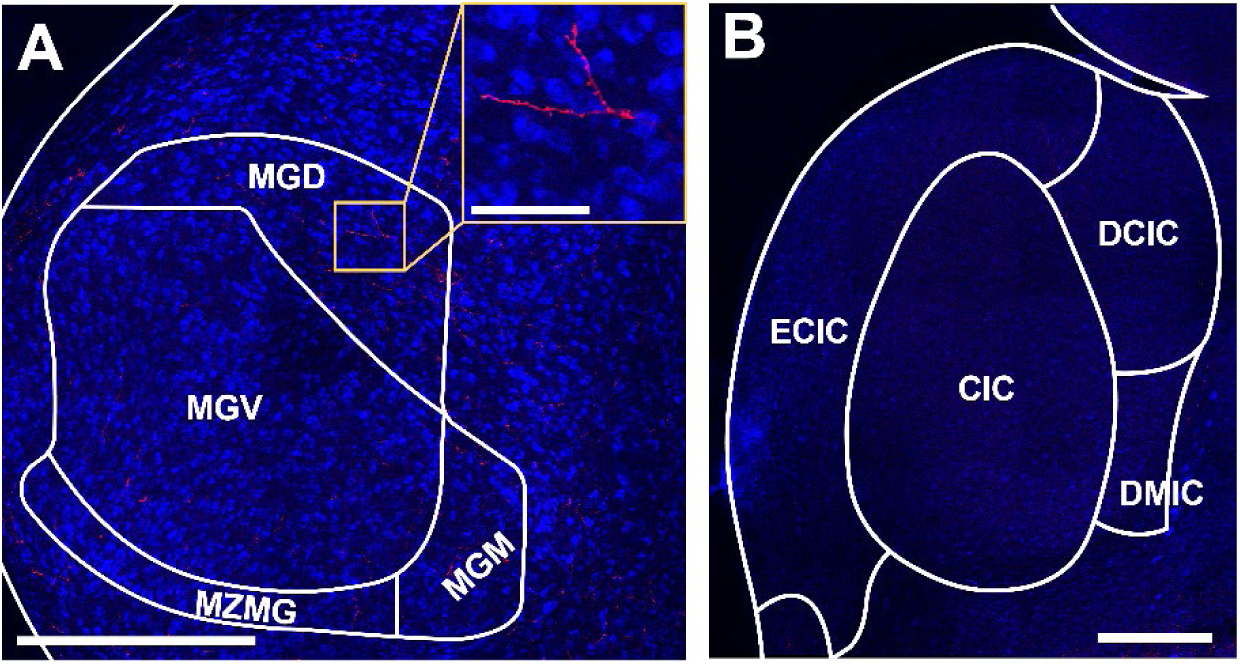
Anterograde tracer into OFC reveals sparse fiber labeling in MGN. **A.** Sparse labeling observed in the MGN in subjects with anterograde tracers injected into the OFC. Scale bar = 500 μm, inset = 100 μm. **B.** No labeling was observed in the IC. Scale bar = 500 μm.

### Auditory cortical-projecting neurons in the OFC send axon collateral to other brain regions

Individual frontal cortical neurons often project to multiple downstream targets, consistent with their role in coordinating brain-wide activity (Gao et al., 2022). To gain insight into how the OFC might specifically orchestrate sound-guided behavior, we asked whether Au1-projecting neurons in the OFC also send axon collaterals to other brain regions. To do so, we injected left Au1 with a virus to drive retrograde expression of Cre-recombinase (AAVrg-hSyn-Cre) and injected left OFC with a virus to drive Cre-dependent expression of mCherry (AAV1-hSyn-DIO-mCherry) in two animals (see Table 1). This intersectional viral strategy selectively labels just the OFC neurons that project to Au1, along with their axon collaterals. We then used tissue clearing and lightsheet microscopy to look for the presence of mCherry fluorescence throughout the intact gerbil brain. In both injected subjects, we observed a dense cluster of brightly labeled cell bodies localized to the ipsilateral OFC (primarily VO and LO; Figure 8A, B), and three distinct axon tracts emanating from these neurons (Figure 8C-F).

**Figure 8.**
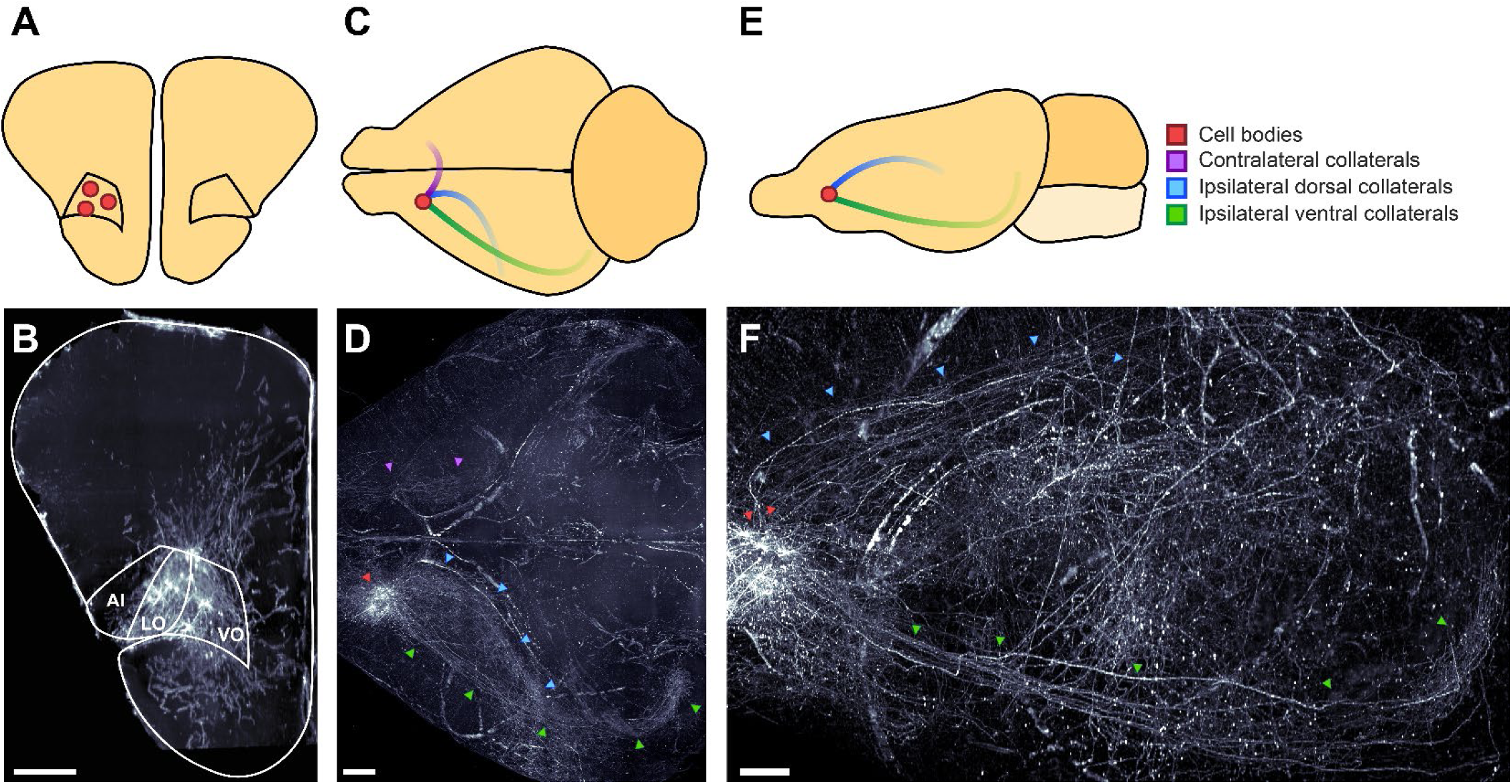
OFC neurons that project to auditory cortex send axon collaterals to other brain regions. **A-B.** Coronal schematic (**A**) and section (**B**) depicting location of identified cell bodies in a representative subject. Section is a 175 μm thick coronal z-stack. Scale bar = 500 μm. **C-D.** Horizontal schematic (**C**) and view (**D**) depicting notable axon tracts emanating from OFC neurons in a representative subject. Scale bar = 1000 μm. **E-F.** Sagittal view (**E**) and section (**F**) depicting axon tracts in representative subject. Scale bar = 1000 μm.

One tract extended dorsally, through the ipsilateral caudate putamen (Figure 9A,B) and through the external capsule to various cortical regions, including the cingulate cortex (Figure 9A,C), somatosensory cortex (Figure 9A,D), auditory cortex (Figure 9E, F), and visual cortex (Figure 9G, H). Within these regions, collaterals targeted multiple cortical layers, and higher-magnification imaging revealed en passant boutons in all regions and terminal boutons in auditory cortex and visual cortex, indicating probable synapses. Additional areas of notable fiber labeling stemming from this axon tract include prelimbic cortex, retrosplenial cortex, and parietal cortex, although we did not confirm the presence of boutons in these regions.

**Figure 9.**
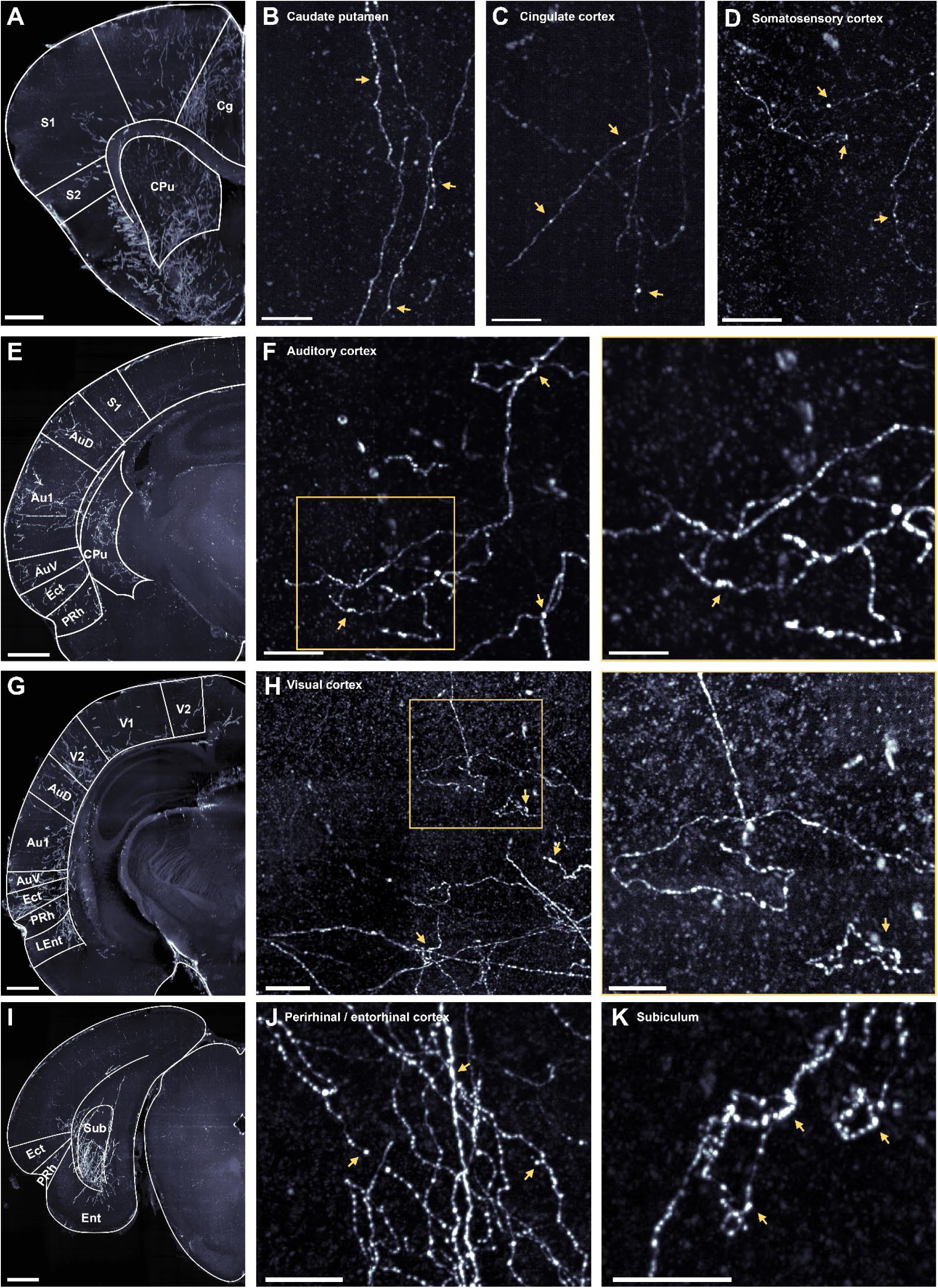
OFC neurons that project to auditory cortex exhibit synaptic boutons in many non-auditory brain regions. **A-D.** Low magnification (**A**) and high magnification (**B-D**) images depicting labeled axon fibers in the anterior caudate putamen (**B**), cingulate cortex (**C**) and the somatosensory cortex (**D**). In B-D, scale bar = 100 μm. Here, and in all following panels, yellow arrows highlight likely synaptic boutons. **E.** Low magnification image depicting labeled axon fibers in the anterior auditory cortex, parahippocampal region, and caudate putamen. **F.** High magnification images depicting labeled axon fibers in the auditory cortex. Scale bars = 100 μm (left), and 50 μm (right). **G.** Low magnification image depicting labeled axon fibers in visual cortex, posterior auditory cortex, and the parahippocampal region. **H.** High magnification images depicting labeled axon fibers in the visual cortex. Scale bars = 100 μm (left), and 50 μm (right). **I.** Low magnification image depicting labeled axon fibers in the subiculum translational area. **J-K.** High magnification images depicting labeled axons fibers in the perirhinal and entorhinal cortices (**J**) and the subiculum (**K**). Scale bars = 50 μm. In panels A, E, G, and I, images are 175 μm thick coronal z-stacks with 1000 μm scale bars.

A second axon tract traveled ventrally to the ipsilateral parahippocampal region (Figure 8C-F). Fibers were observed in the ectorhinal, perirhinal, and entorhinal cortices (Figure 9E, G, I, J), and particularly dense labeling was observed in the subiculum (Figure 9I, K). These fibers exhibited en passant and terminal boutons, suggesting the presence of functional synapses.

A final axon tract crossed the midline and then branched extensively, targeting the contralateral OFC, cingulate cortex, caudate putamen, auditory cortex, and parahippocampal region (Figure 8C-D). A complete list of regions where we observed labeled fibers, along with a qualitative estimate of the intensity of labeling is provided in Table 7.

**Table 7.**
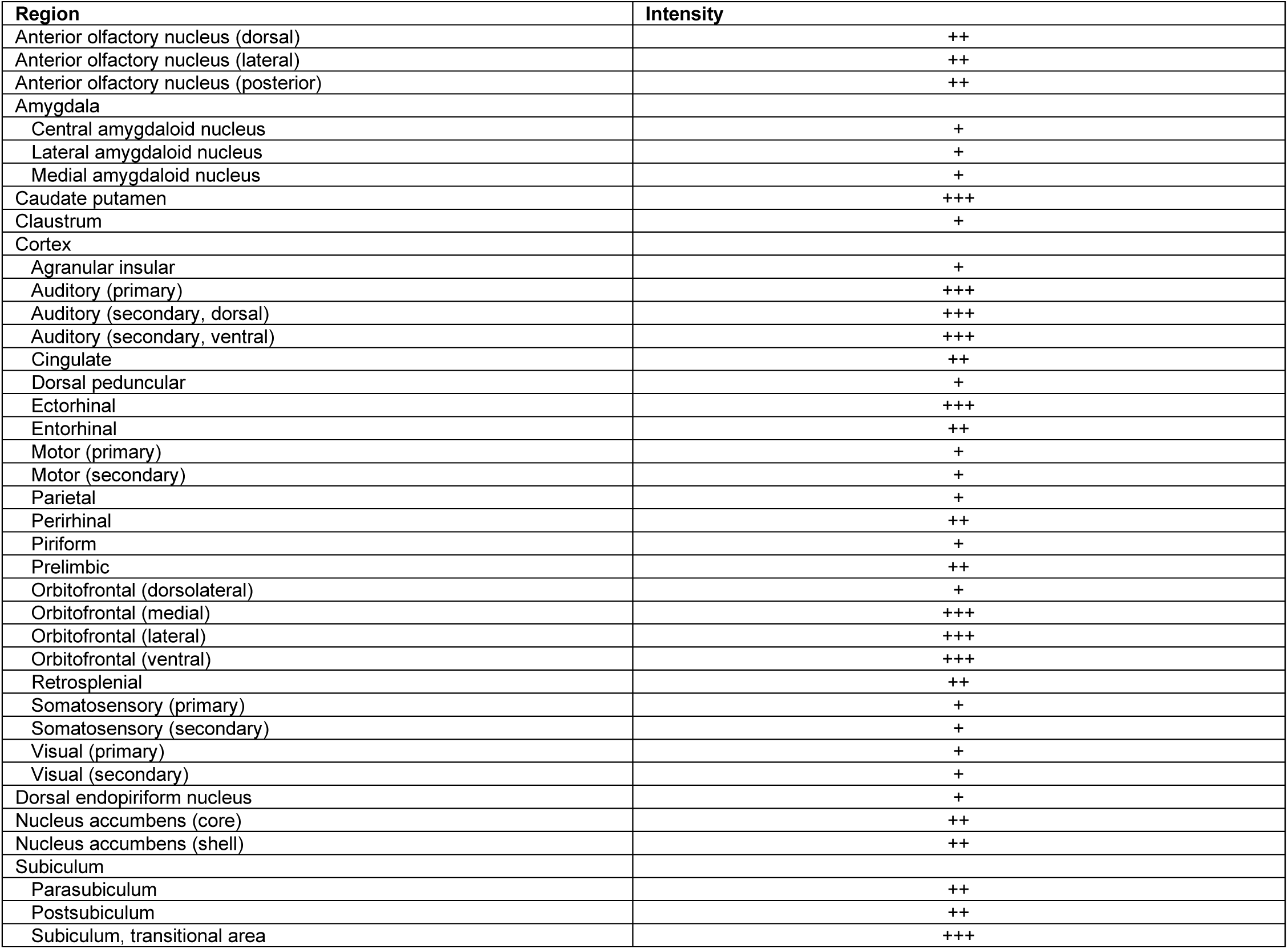
Qualitative estimate of the intensity of labeled fibers in various brain regions innervated by OFC-Au1 axon collaterals.

## Discussion

Using anterograde and retrograde AAV tracers, we characterized the descending projections from the OFC to cortical and subcortical auditory regions in an established model organism for auditory research, the Mongolian gerbil. We found that the OFC exhibits little to no connectivity with the MGN and IC, but robustly innervates auditory cortices. The pattern of connectivity varied across OFC’s mediolateral axis, such that nearly all auditory cortical-projecting neurons were observed in the ventral and medial subdivisions. These neurons preferentially targeted infragranular layers, with the densest innervation in secondary auditory cortices. An intersectional viral strategy, tissue clearing, and lightsheet microscopy further revealed that the OFC neurons that project to the auditory cortex also send wide-ranging axon collaterals to distant targets, including many non-sensory areas. Our results provide an anatomical basis for understanding the role of the OFC in shaping auditory function and sound-guided behavior.

In Au1, we found OFC axons in all cortical layers, suggesting that OFC is well-positioned to directly shape sensory-evoked responses throughout the cortical column. Indeed, optogenetic stimulation of OFC axon terminals in mice activates multiple Au1 laminae and alters L2/3 receptive fields (Winkowski et al., 2018). Our data suggest that OFC’s influence on Au1 sound processing may be largely mediated via infragranular activation, as OFC fiber innervation is strongest in L5/6. Neurons in L5/6 provide extensive feedback to subcortical auditory structures (for review see Winer, 2006), and make local connections within Au1 (Guo et al., 2017; Onodera & Kato, 2022; Prieto & Winer, 1999; Williamson & Polley, 2019). These pathways shape spectrotemporal tuning and response gain in the inferior colliculus, medial geniculate body, and auditory cortex (Antunes & Malmierca, 2021; Gao & Suga, 1998; Ma & Suga, 2001; Malmierca et al., 2015; Saldeitis et al., 2021; Zhang & Yan, 2008), and directly impact behavioral performance on auditory detection and discrimination tasks (Guo et al., 2017; Homma et al., 2017). In addition, L6 corticothalamic circuits have been hypothesized to gate the transmission of behaviorally-relevant information from the periphery to the cortex (He, 1997; Sherman & Koch, 1986; Yu et al., 2004; Zhang et al., 1997). Taken together, these findings suggest that OFC may mediate rapid changes in stimulus salience by activating both intracortical and subcortical feedback projections originating in the deep layers of auditory cortex. Future experiments examining OFC’s impact on auditory cortical infragranular activity *in vivo* are needed, as prior reports have exclusively monitored and/or manipulated OFC axon terminals in L2/3 (Liu et al., 2021; Winkowski et al., 2018).

Sound-evoked representations are transformed as they traverse the auditory cortical hierarchy, gradually incorporating information about stimulus meaning or emotional valence as the signals move from primary to secondary and tertiary regions (Atiani et al., 2014; Elgueda et al., 2019; Grosso et al., 2015; Takahashi et al., 2010, 2011; Yin et al., 2020). Our finding that OFC sends a larger projection to secondary auditory cortical regions than to Au1 suggests one anatomical basis for these observations. Higher-order auditory regions also exhibit stimulus-evoked activity that persists for hundreds of milliseconds after sound offset (Atiani et al., 2014; Elgueda et al., 2019) and long temporal integration windows (Bendor & Wang, 2007; Boemio et al., 2005; Scott et al., 2011), both of which offer an increased opportunity for signals encoding behavioral outcomes to be combined with sensory input. These findings, coupled with VO’s proposed role in signaling reward history, prediction error, and value (Riceberg & Shapiro, 2017; Steiner & Redish, 2012; Stolyarova & Izquierdo, 2017; Young & Shapiro, 2011), suggest that OFC’s robust projection to secondary auditory cortices may serve to rapidly update sensory representations during sound-guided behavior. Recent work suggests that OFC inputs to primary sensory cortical regions may also serve a similar function (Banerjee et al., 2020; Liu et al., 2020; Liu et al., 2021).

We found that auditory cortical-projecting OFC neurons also send wide-ranging axon collaterals to various brain regions involved in learning, memory, and decision making. One downstream target that receives substantial collateral innervation is a striatal area strongly implicated in auditory behavior—the posterior portion of the caudate putamen (CPu). The posterior CPu receives direct projections from Au1 (Budinger et al., 2000a, 2008; Roger & Arnault, 1989), and these inputs support associative learning by integrating information about sound identity, behavioral choice, and expectation of reward (Guo et al., 2018, 2019; Xiong et al., 2015; Znamenskiy & Zador, 2013). These findings have led some to hypothesize that many of the response properties exhibited by the posterior CPu are inherited directly from Au1 (Guo et al., 2019). Our results suggest an alternative, though not mutually exclusive, model in which Au1 and posterior CPu exhibit similar responses in part because they receive input from the same OFC neurons. It is also interesting to note that within the CPu, the pattern of Au1 innervation matches the pattern of innervation from axon collaterals branching from the OFC to Au1 projection (Budinger et al., 2000b; Roger & Arnault, 1989). Taken together, these results suggest that Au1 and OFC may work in tandem to generate stable striatal representations of learned stimulus values during sound-guided behavior.

We also observed a significant collateral projection to the parahippocampal region, which is comprised of entorhinal, ectorhinal, and perirhinal cortices, as well as the subiculum. These structures are critical for working memory (Nemanic et al., 2004; Young et al., 1997). Though the anatomical connection between the OFC and the parahippocampal region is well documented (Burwell & Amaral, 1998; Hoover & Vertes, 2011; Kondo & Witter, 2014; Witter et al., 2017), our understanding of the function of this pathway is limited. OFC, sensory cortices, and the parahippocampal region all display stimulus-selective activity when sensory stimuli are held in working memory (Gottlieb et al., 1989; Ramus & Eichenbaum, 2000; Young et al., 1997). Our observation that the same OFC neurons that project to the auditory cortex also project to the parahippocampal region suggests that these neurons are particularly well-positioned to coordinate activity between these two areas during working memory. Future experiments employing targeted manipulations of OFC axon terminals will be needed to reveal whether these pathways do indeed form part of a sensory mnemonic circuit.

Collectively, our data reveal a long-range projection from the OFC that is particularly well-positioned to rapidly update stimulus representations in the gerbil auditory cortex and orchestrate brain-wide changes during auditory learning. These findings advance our understanding of the neural circuits that support auditory cognition and perceptual flexibility.

## Author contributions

M.L.C, L.H., and R.Y. designed the research; R.Y., L.H., L.N., R.B., and N.M. performed the research; R.Y. and M.L.C. wrote the paper.

## Acknowledgements

This work was supported by National Institute of Health Grant K99/R00DC016046 to M.L.C and T32DC00046 to R.Y. Purchase of the Zeiss LSM 980 Airyscan 2 was supported by Award Number 1S10OD025223-01A1 from the National Institute of Health. We thank Dr. Dan Sanes (New York University) for the use of his lab for one of the reported procedures. We also thank Dr. Daniel Stolzberg (University of Maryland) for help with image processing, Dr. Daniel Miller (University of Illinois at Urbana-Champaign) for advice on data analysis, Dr. Matheus Macedo-Lima (University of Maryland) for help with statistics, and all members of the M.L.C. laboratory for their constructive criticism and support. The authors declare no competing financial interests.

## Abbreviations

AAF: Anterior auditory field
AuD: Dorsal auditory cortex
AuV: Ventral auditory cortex
Au1: Primary auditory cortex
CIC: Central nucleus of the inferior colliculus
Cg: Cingulate cortex
CPu: Caudate putamen (striatum)
DCIC: Dorsal cortex of the inferior colliculus
DLO/AI: Dorsolateral orbitofrontal cortex/agranular insular cortex
DMIC: Dorsomedial nucleus of the inferior colliculus
ECIC: External cortex of the inferior colliculus
Ect: Ectorhinal cortex
Ent: Entorhinal cortex
IC: Inferior colliculus
LO: Lateral orbitofrontal cortex
PRh: Perirhinal cortex
MGN: Medial geniculate nucleus
MGD: Medial geniculate nucleus, dorsal part
MGM: Medial geniculate nucleus, medial part
MGV: Medial geniculate nucleus, ventral part
MZMG: Marginal zone of the medial geniculate
MO: Medial orbitofrontal cortex
Sub: Subiculum translational area
S1: Primary somatosensory cortex
S2: Secondary somatosensory cortex
OFC: Orbitofrontal cortex
VO: Ventral orbitofrontal cortex
V1: Primary visual cortex
V2: Secondary visual cortex

## Notes

### Competing Interest Statement

The authors have declared no competing interest.

